# Regionalized tissue fluidization by an actomyosin cable is required for epithelial gap closure during insect gastrulation

**DOI:** 10.1101/744193

**Authors:** A. Jain, V. Ulman, A. Mukherjee, M. Prakash, L. Pimpale, S. Muenster, R. Haase, K.A. Panfilio, F. Jug, S.W. Grill, P. Tomancak, A. Pavlopoulos

**Affiliations:** Max-Planck-Institute of Molecular Cell Biology and Genetics, Dresden, Germany; Technische Universität Dresden, Dresden, Germany; IT4Innovations, Technical University of Ostrava, Czech Republic; Max-Planck-Institute for the Physics of Complex Systems, Dresden, Germany; Center for Systems Biology, Dresden, Germany; Biotechnology Center, TU Dresden, Germany; Institute for Zoology: Developmental Biology, University of Cologne, Cologne, Germany; School of Life Sciences, University of Warwick, Coventry, UK; Cluster of Excellence Physics of Life, TU Dresden, Germany; Janelia Research Campus, Howard Hughes Medical Institute, Ashburn, United States; Institute of Molecular Biology and Biotechnology, Foundation for Research and Technology-Hellas, Heraklion, Greece

## Abstract

Many animal embryos pull and close an epithelial sheet around the spherical or ellipsoidal egg surface during a gastrulation process known as epiboly. The ovoidal geometry dictates that the epithelial sheet first expands and subsequently compacts. Moreover, the epithelial sheet spreading over the sphere is mechanically stressed and this stress needs to be released. Here we show that during extraembryonic tissue (serosa) epiboly in the red flour beetle *Tribolium castaneum*, the non-proliferative serosa becomes regionalized into two distinct territories: a dorsal region under higher tension away from the leading edge with larger non-rearranging cells, and a more fluid ventral region under lower tension surrounding the leading edge with smaller cells undergoing cell intercalation. Our results suggest that fluidization of the leading edge is caused by a heterogeneous actomyosin cable that drives sequential eviction and intercalation of individual cells away from the serosa margin. Since this developmental solution utilized during epiboly resembles the mechanism of wound healing in other systems, we propose actomyosin cable-driven local tissue fluidization as a conserved morphogenetic module for closure of epithelial gaps.

---

Epiboly is one of the evolutionarily conserved morphogenetic movements during animal gastrulation^1^. It involves spreading of an epithelial sheet over the spherical or ellipsoidal egg. The sheet eventually forms a continuous layer that entirely surrounds the embryo and the yolk sac. During this morphogenetic event, fundamental geometrical and mechanical problems arise. First, in order to cover the entire egg, the epithelium has to expand in surface area. However, once the egg equator is reached, the expanding tissue must also undergo a regional compaction at its leading edge in order to seal seamlessly at the bottom of the sphere (**Fig 1A**). Studies in fish showed that the tissue spreading is mediated by changes in cell shape, cell number and cell arrangement coupled to constriction of an actomyosin ring in the yolk at the leading edge of the sheet^2–5^. However, it remains unclear whether pulling forces at the leading edge expand cells uniformly throughout the tissue and how cells behave at the leading edge that needs to compact. Second, spreading over a sphere induces mechanical stress in the tissue. In zebrafish, mechanical stress during epibolic expansion is released by oriented cell divisions in the tissue^4^. In other epiboly systems however, cell division does not occur, and thus other unknown mechanisms have to alleviate built-up stress.

**Figure 1:**
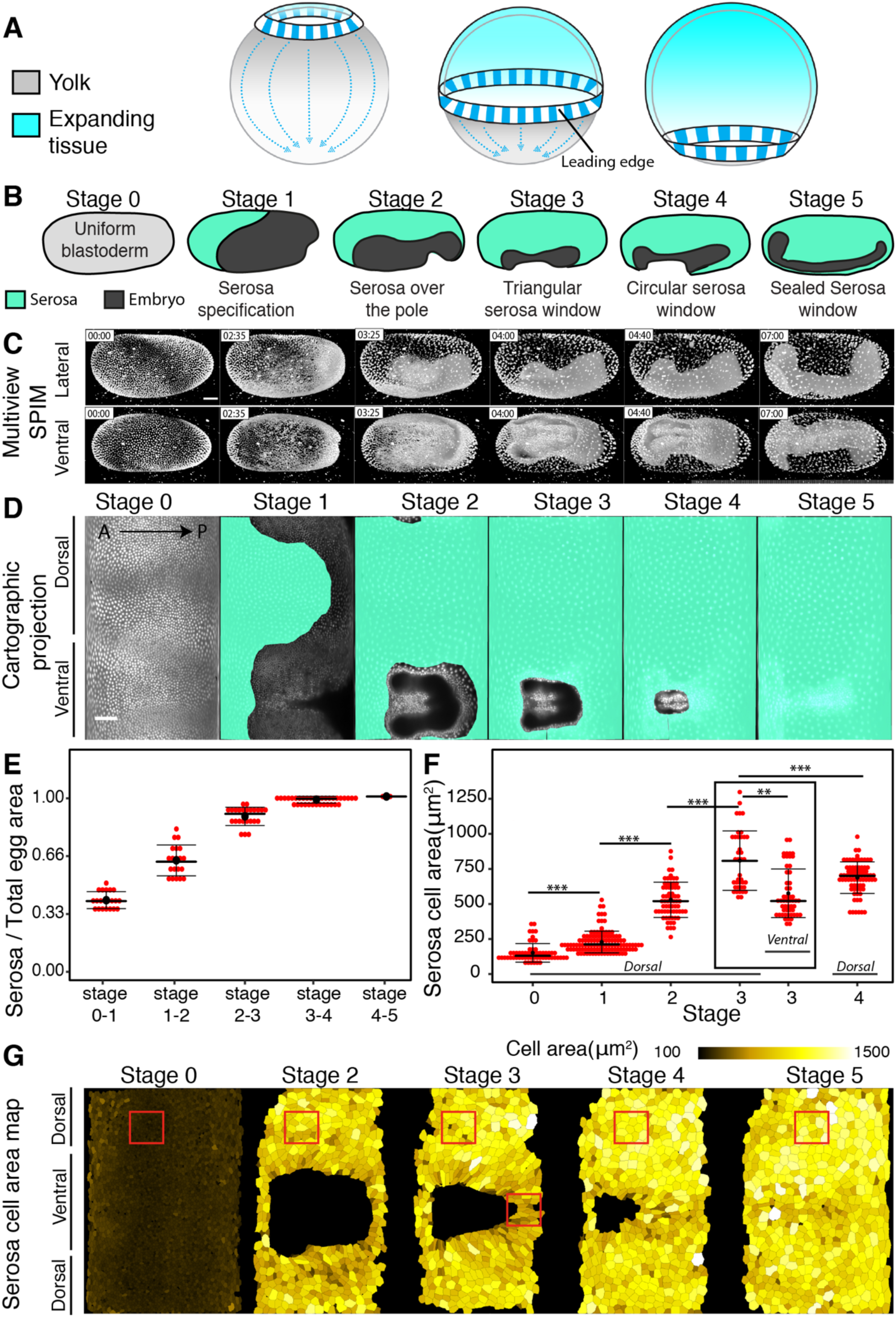
Inhomogeneous tissue expansion during *Tribolium* serosa morphogenesis. **(A)** Schematic depiction of the geometric constraints experienced by a tissue expanding over a spherical yolk cell. The leading edge undergoes an area increase followed by an area decrease after it crosses the equator. **(B)** Illustrations of the stages of *Tribolium* embryogenesis from cellular blastoderm to serosa window closure. **(C)** 3D rendered reconstructed multi-view time-lapse SPIM recording of a *Tribolium* embryo expressing the fluorescent EFA:nGFP nuclear marker. The embryo is shown from the lateral and ventral view at the 6 reference stages corresponding to the schematics in (B). All imaged embryos in this and other panels are shown with anterior to the left and all time stamps are in hours: minutes. Scale bar is 50 *µ*m. **(D)** 2D cartographic projection of a 4D SPIM recording of *Tribolium* embryo expressing EFA:nGFP. The extent of the serosa tissue is highlighted in turquoise. Scale bar is approximately (see Methods) 100 *µ*m. **(E)** The area of the serosa tissue calculated from cartographic projections of 4D SPIM recordings. The data are normalized to the total serosa area at stage 5 in each case. For every stage the total serosa area is calculated for all timepoints between two consecutive stages in three different embryos and plotted as a distribution. Plots in this and all other panels indicate the median with a thick line, the mean with a black dot and the standard deviation with the thin error bars **(F)** Comparison of the distributions of apical areas of cells sampled from point scanning confocal recordings of *Tribolium* embryos expressing LifeAct-eGFP membrane marker at reference stages labeled according to (B). The number of cells (n) and the number of embryos (N) sampled at different stages were as follows: In the dorsal region, Stage 0 n=58, N=6, Stage 1 n=116, N=11, Stage2 n=66, N=9, Stage 3 n=39, N=6, Stage 4 n=76, N=10 and Stage 3 ventral n= 52, N=7. The difference between distributions was tested using unpaired Welch’s t-tests (same for all Figures unless stated otherwise). P-values between 0.05-0.01 are labelled with *, 0.009-0.001 are labeled with **, <0.001 with *** and ns signify non-significant p-values (same for all Figures). **(G)** Cartographic projections of reference stages of an embryo labelled with LifeAct-eGFP and imaged live with multi-view SPIM. The projections are overlaid with manually curated automated segmentation results visualizing apical areas of serosa cells through a color code. Red boxes indicate the approximate regions from which cells were sampled in confocal datasets quantified in (F).

For example, in many insect taxa, the developing embryo is completely surrounded by a protective epithelial cell layer of extraembryonic fate called the serosa^6–10^. In the red flour beetle, *Tribolium castaneum*, extraembryonic serosal cells are initially specified as an anterior cap of the cellular blastoderm, which subsequently spreads over the gastrulating embryonic part of the blastoderm^11^. This process resembles vertebrate epiboly but occurs in the complete absence of serosal cell division (**Fig 1B**). The spreading serosal tissue expands over the posterior pole and eventually closes ventrally over the contracting embryo in a process known as serosa window closure^12–14^. However, it is not understood how the leading serosal cells at the rim of the serosa window achieve final compaction. It is also unknown if and how mechanical tension arises and gets released in the serosal tissue during spreading.

To address these questions, we used the *Tribolium* serosal epiboly and closure as a model to understand how the mechanical properties of cells and physical forces are regionalized to wrap a non-dividing epithelial sheet around an ellipsoidal egg. We imaged the progression of serosa spreading with multi-view light sheet microscopy in embryos expressing a nuclei-marking eGFP (**Fig 1C, *Supplementary movie 1***). Taking advantage of the serosa’s topology as a superficial egg layer, we unwrapped the 3D data into 2D cartographic time-lapse projections and segmented the serosal part of the blastoderm tissue^15^ (**Fig 1D, *Supplementary Fig 1A-D, Supplementary movie 2*** and ***11***). The serosa covered initially about 35% of the egg surface and spread to cover 100% of the surface (**Fig 1E**). In order to examine the expansion at the cellular level, we imaged embryos expressing LifeAct-eGFP that labels cortical F-actin^13,16^ and segmented the apical surface of all serosal cells at 5 reference stages (**Fig 1B**) during serosal expansion (**Fig 1G**). The results showed that the ∼ 3-fold expansion in serosal tissue surface area was mirrored by a ∼ 3-fold expansion of the apical area of serosal cells from stage 1 to stage 4 (**Fig 1F)**. Strikingly, serosal cells did not expand uniformly: at stage 3, the apical area of ventral cells in the vicinity of the serosa window were on average 29% smaller compared to dorsal cells (**Fig 1F-G**). We conclude that serosal epiboly exhibits inhomogeneous apical cell area expansion in order to accommodate the ventral area compaction required by the elliptical geometry of the egg.

An alternative but not mutually exclusive mechanism to achieve ventral area compaction is by reducing the number of marginal cells at the serosa window (**Fig 2A**)^12^. While it is in principle possible that leading cells are not excluded and converge to a multicellular rosette, such a rosette has not been observed during *Tribolium* serosa window closure^12,13,17^. Our cell tracking experiments showed that the initial number of approximately 75 leading cells progressively decreased to only 5-6 cells during final serosal closure (**Fig 2B,C**) and that these cells originated from all around the periphery of the window (**Fig 2D, *Supplementary movie 3***). Careful examination of individual cells at the leading edge in time-lapse recordings of embryos of the LifeAct-eGFP transgenic line revealed frequent rearrangement of cells resulting in cells leaving the serosa edge (**Fig 2E,F, *Supplementary movie 4***). The leaving cells shrunk their leading edge facing the serosa window and elongated radially in the direction approximately orthogonal to the window (***Supplementary Figure 2A-C***). Upon leaving the edge, the cells gradually relaxed to a hexagonal shape as they reintegrated into the bulk of the tissue (***Supplementary Fig 2D***). Mapping of those behaviours onto the time-lapse cartographic projections revealed that the serosa was regionalized into two distinct territories. Dorsal cells, several cell diameters away from the edge, were hexagonally packed, isotropically stretched and showed no significant neighbor exchanges. By contrast, ventral cells surrounding the serosa window were irregularly packed, showed anisotropically stretched shapes (***Supplementary Fig 3A***) and frequently exchanged neighbors (**Fig 2H, *Supplementary movie 5*** and ***11***).

**Figure 2:**
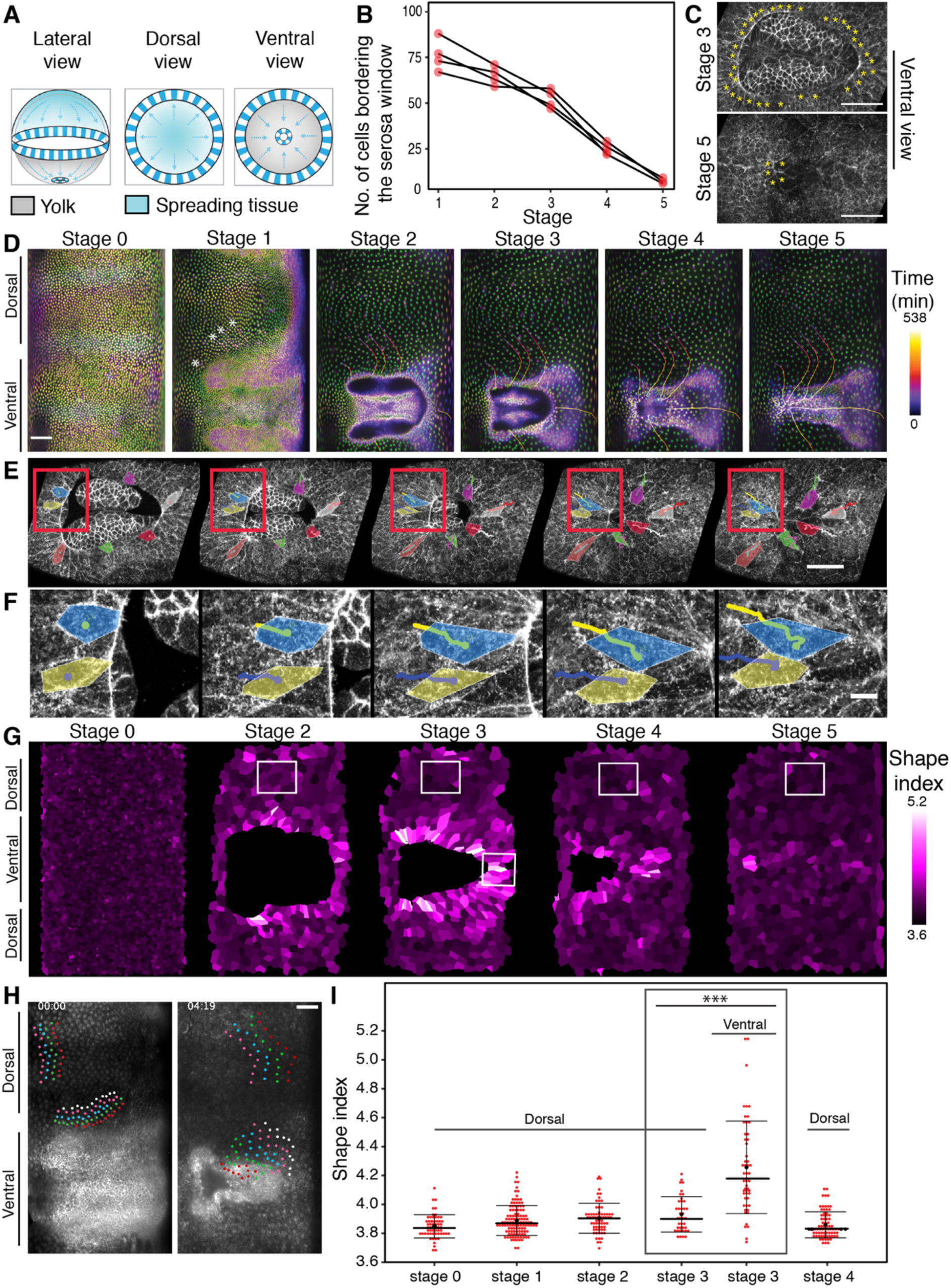
Cell behaviors at the ventral leading edge of the serosa window are distinct from the behaviors in dorsal serosa. **(A)** Schematic illustration of the putative mechanism of closing serosa window by reducing the number of cells at the leading edge of the window over time. **(B)** Plot of the total number of cells at the embryo-serosa boundary during serosa window closure counted at the five reference stages. (N=4) **(C)** Confocal images highlighting the cells (yellow asterisks) forming the leading edge of the serosa window at Stage 3 (top) and Stage 5 (bottom). Scale bar is 10 *µ*m. **(D)** Cartographic projection of a Histone-eGFP labelled embryo imaged with multi-view SPIM. The progressively deeper onion layers of the projection are color-coded to distinguish superficial and internal nuclei. The nuclei participating in closing of the serosa window were back-tracked to the uniform blastoderm stage to reveal their spatial origin. Tracks are color-coded by time as indicated by the color scale. Scale bar is approximately 100 *µ*m. **(E)** Frames from a confocal recording of the serosa window closure in embryos expressing LifeAct-eGFP. Selected cells at the leading edge of the serosa window are outlined and colored to show that some cells shrink their serosa-window-facing membranes and planarly intercalate into the serosa epithelium. Red box marks the inset shown in (**F**). Scale bar is 50 *µ*m in (E) and 10 *µ*m in (F). **(G)** Cartographic projections of an embryo labelled with LifeAct-eGFP, imaged in SPIM and semi-automatically segmented. The color code indicates the value of the shape index for each segmented serosa cell. White boxes indicate the approximate regions from which cells were sampled in confocal datasets quantified in (I). **(H)** Cartographic projections of an embryo labelled with LifeAct-eGFP at the beginning (left) and towards the end (right) of serosa window closure. Selected cells were tracked over time and color coded to visualize the difference in the extent of neighbor exchanges between the dorsal cells and ventral cells close to the leading edge of the serosa widow. Scale bar is approximately 100 *µ*m. **(I)** Distributions of shape indices of cells segmented from Stage 0-4 LifeAct-eGFP embryos imaged by point scanning confocal. Numbers of cells and embryos are the same as in 1F.

Movement of cells past each other during neighbor exchange has been linked to increased tissue fluidity^18–21^. A useful theoretical framework to assess the behavior of the serosal tissue is the shape index analysis that infers solid-like or fluid-like tissue states from cell shapes in epithelia^22–24^. The theory predicts a critical value of shape index *p* = 3.81 marking the transition from a solid-like to a fluid-like behaviour (but see also^25^ and below). Our results showed that at stage 3 ventral cells had on average a high shape index 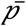 of 4.25 characteristic of fluid-like tissues, unlike dorsal cells that had a significantly lower 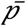 value of 3.93, which indicated that the ventral region is much more fluid-like compared to the dorsal region (**Fig 2G,I, *Supplementary Movie 11***). These results raised the hypothesis that during serosal epiboly the tissue in the vicinity of the window undergoes a solid-to-fluid structural transition (fluidization) that unjams the tissue and enables seamless closure.

We next asked what the mechanical function of the ventral serosa fluidization could be. If the dorsal serosa behaves as a solid-like material, we expect that while being pulled over the egg it would increasingly build up tension. This rising tension would make it increasingly more difficult to further close the serosa window. The function of the ventral cell rearrangement in the proximity of the serosa window could then be in releasing this tension to facilitate closure. Consequently, we would predict a difference in tissue tension between dorsal and ventral serosa. To test this, we performed laser ablations, inflicting large incisions across 3-4 cells at different reference stages and positions and compared the recoil velocities^26^ (**Fig 3A,B**). These tissue cutting experiments showed that the tension in the dorsal side increased progressively as the serosa expanded posteriorly and ventrally around the posterior pole and plateaued after the serosa window formed (**Fig 3C**). The intact cells neighboring the ablation site responded to the release in tissue tension post-ablation by immediately decreasing their apical areas by 1/3^rd^ ***(Supplementary Fig 4).*** These two results suggest that the dorsal tissue behaves as an elastic solid from a mechanical perspective. Importantly, for incisions that were inflicted at the ventral side of the serosa exhibiting cell rearrangements, the tension was lower compared to the dorsal side (**Fig 3D**). Thus, laser cutting experiments corroborate the regionalization of the serosa into a more solid dorsal region that stays under high tension and a more fluid ventral region that has relaxed its tension.

**Figure 3:**
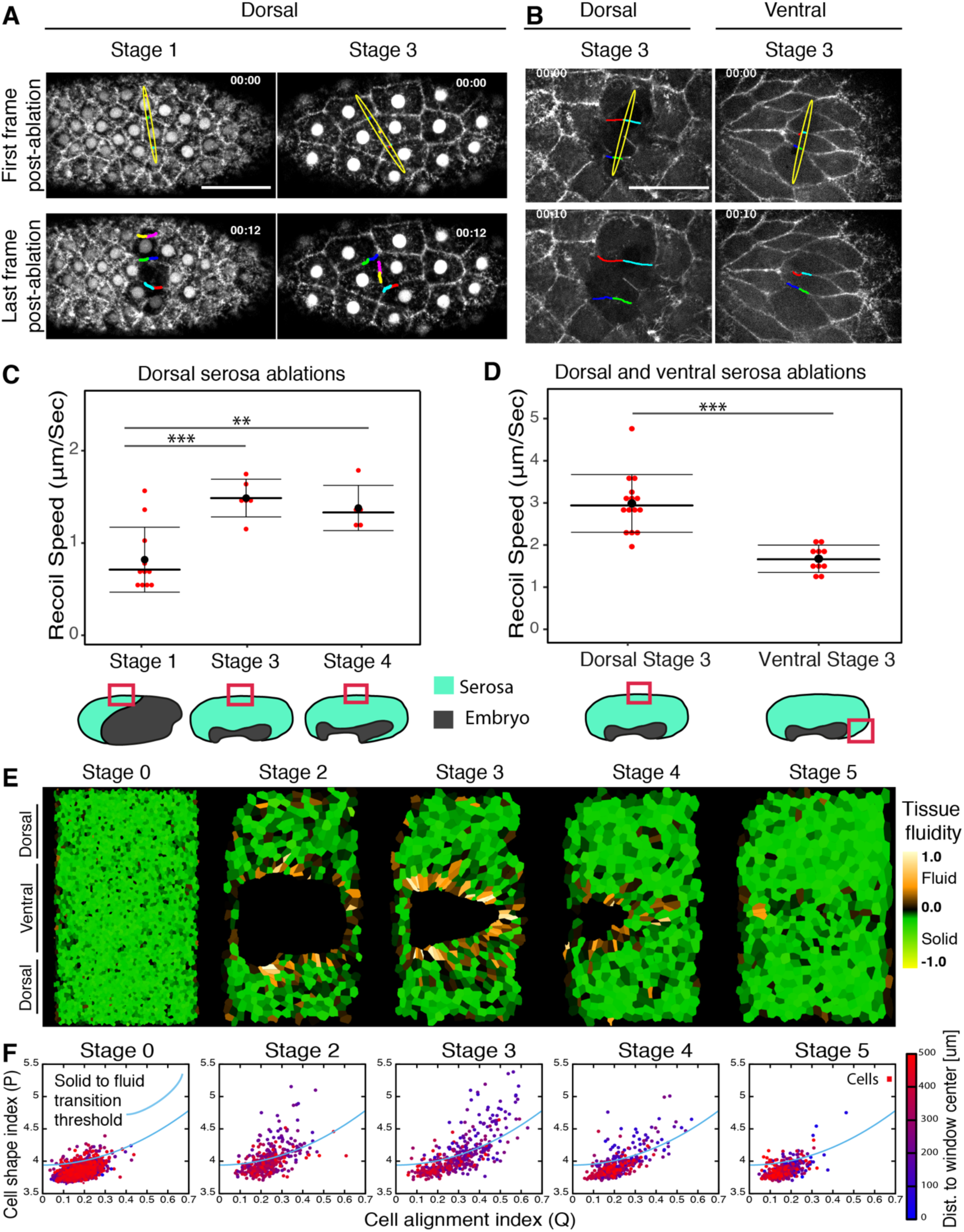
Tension landscape in the expanding serosa. **(A)** Tissue laser ablations in the dorsal serosa at different reference stages. Images show the serosal tissue before (top) and after (bottom) laser ablation in the dorsal region of the embryo expressing LifeAct-eGFP and EFA:nGFP at Stage 1 and Stage 3. The yellow ellipses show the extent of the cut. The colored lines highlight the displacement of the severed cell edges. Scale bar is 50 *µ*m. **(B)** Comparison of laser ablation in dorsal and ventral regions of the serosa (as depicted schematically below the graph in (D)) at stage 3 in distinct embryos expressing LifeAct-eGFP. The yellow ellipse shows the extent of the cut. Scale bar is 50 *µ*m. **(C)** Graph shows comparison of recoil velocities after laser ablation at Stages 1, 3 and 4. The ablations were performed at the same dorsal position of each embryo as indicated by the red box in the reference stage schematics below. Each dot represents one cut in one embryo. The number of embryos (N) sampled at different stages were as follows: Stage 1 N=11, Stage 3 N = 6, Stage 4 N = 5. **(D)** Graph shows comparison of recoil velocities after laser ablation of serosal cells in the dorsal and the ventral regions of Stage 3 embryos. The red boxes in the embryo illustrations below the graphs indicate the position of the dorsal and ventral cuts for the data shown in (B) and (D). Each dot represents one cut in one embryo. The number of embryos (N) sampled were as follows: Dorsal N=15, Ventral N = 10. **(E)** Cartographic projections of an embryo labelled with LifeAct-eGFP, imaged in SPIM and semi-automatically segmented. The color code indicates the tissue fluidity measured by subtracting the local solid-to-fluid transition shape index threshold (blue curve in (F)) from the cell shape index for each segmented cell (see methods). Positive values indicate fluid-like tissue while negative values indicate a solid-like tissue. **(F)** Scatter plot of cell shape alignment index (x-axis) and shape index (y-axis) values of individual cells in the maps shown in (E). The cells are color coded according to their distance from the center of the serosa window. The blue line indicates theoretically predicted threshold value of shape index signifying solid-to-fluid structural transition. Points below the line indicate solid-like cells and points above the line fluid-like cells.

While the tension profile supports the hypothesis of ventral tissue fluidization suggested by the shape index analysis, it had been recently shown that the relationship between shape index and tissue fluidity is non-linear when a tissue is under anisotropic tension^25^. Since we obtained evidence that the *Tribolium* serosa exhibits anisotropic tension, we applied this extended theoretical framework, and calculated a local cell alignment factor *Q* across the serosal tissue (see Methods)^27^. The theory predicts that for a given value of *Q* the shape index *p* needs to exceed an adjusted threshold value in order for the tissue to be fluid-like. For each local value of *Q* across the cartographic maps, we plotted the difference between the actual shape index value (*p)* of the cell and the local threshold signifying solid-to-fluid transition (**Fig 3E,F, *Supplementary Movie 11***). This analysis revealed that also when taking tissue tension anisotropy into consideration, the ventral cells lining the rim of the serosa window exhibited a distinct fluid-like state during closure in stark contrast with the rest of the epithelium exhibiting a solid-like state. Therefore, both experimental and theoretical evidence support the local fluidization of the ventral-most serosal tissue.

We next asked what induces the local tissue fluidization. Recent live imaging studies of *Tribolium* gastrulation suggested that an accumulation of actin, resembling a cable, emerges at the leading edge of the serosa^13,28^. To test whether this accumulation indeed represents a contractile actomyosin cable, we imaged the distribution of non-muscle myosin II (hereafter referred to as myosin) in gastrulating embryos injected with the Tribolium myosin regulatory light chain mRNA fused to eGFP (Tc-sqh-eGFP). During epiboly, myosin accumulated at the boundary between the serosa and the embryonic primordium (**Fig 4A,B, *Supplementary Fig 5A,C, Supplementary movie 6***). Actomyosin enrichment at the serosa-embryonic boundary initiated shortly after epiboly started, and became more pronounced as the boundary stretched around the posterior pole. It peaked during serosa window closure and at this stage appeared as a contiguous supra-cellular cable (**Fig 4B, *Supplementary Fig 5A*,C**). The actomyosin cable lined the rim of the serosa window and underwent shape transformations from triangular to spherical during closure (**Fig 4C, *Supplementary Fig 5B***). By segmenting and measuring the length and intensity of LifeAct accumulation, we found that the cable first increased its length until the serosa-embryonic boundary reached the posterior pole and then decreased in length to zero during window closure (***Supplementary Fig 5D*).** As the cable shrunk, the total myosin intensity normalized by cable length stayed the same or increased over time (***Supplementary Fig 5E***). Laser cutting experiments of individual cell edges contributing to the actomyosin cable revealed that the cable was under tension and that this tension increased over time (**Fig 4D,E,F, *Supplementary movie 7***). If the cable acted as a contiguous contractile ring, one would expect global loss of tension after a cut. Instead, when we inflicted successive laser cuts at different positions of the same cable, the recoil velocities were comparable (**Fig 4G**). This indicated that individual cells of the cable contract their myosin-loaded edges independently and implied that the serosa window edge acts as a chain of independently contractile units. Moreover, the myosin distribution around the cable circumference showed strong heterogeneity, with some cells exhibiting higher and other cells exhibiting lower myosin accumulation. Cells with more myosin contracted their cable-forming edges and were evicted from the leading edge of the serosa earlier than cells with lower levels of myosin (**Fig 4H,I, *Supplementary movie 8***). Since the myosin intensity correlates with the cell leaving behavior, we conclude that differential line tension along the cable circumference drives the eviction of the cells from the cable and the resulting cell rearrangements lead to tissue fluidization and eventual closure of the epithelial gap (**Fig 4J**).

**Figure 4:**
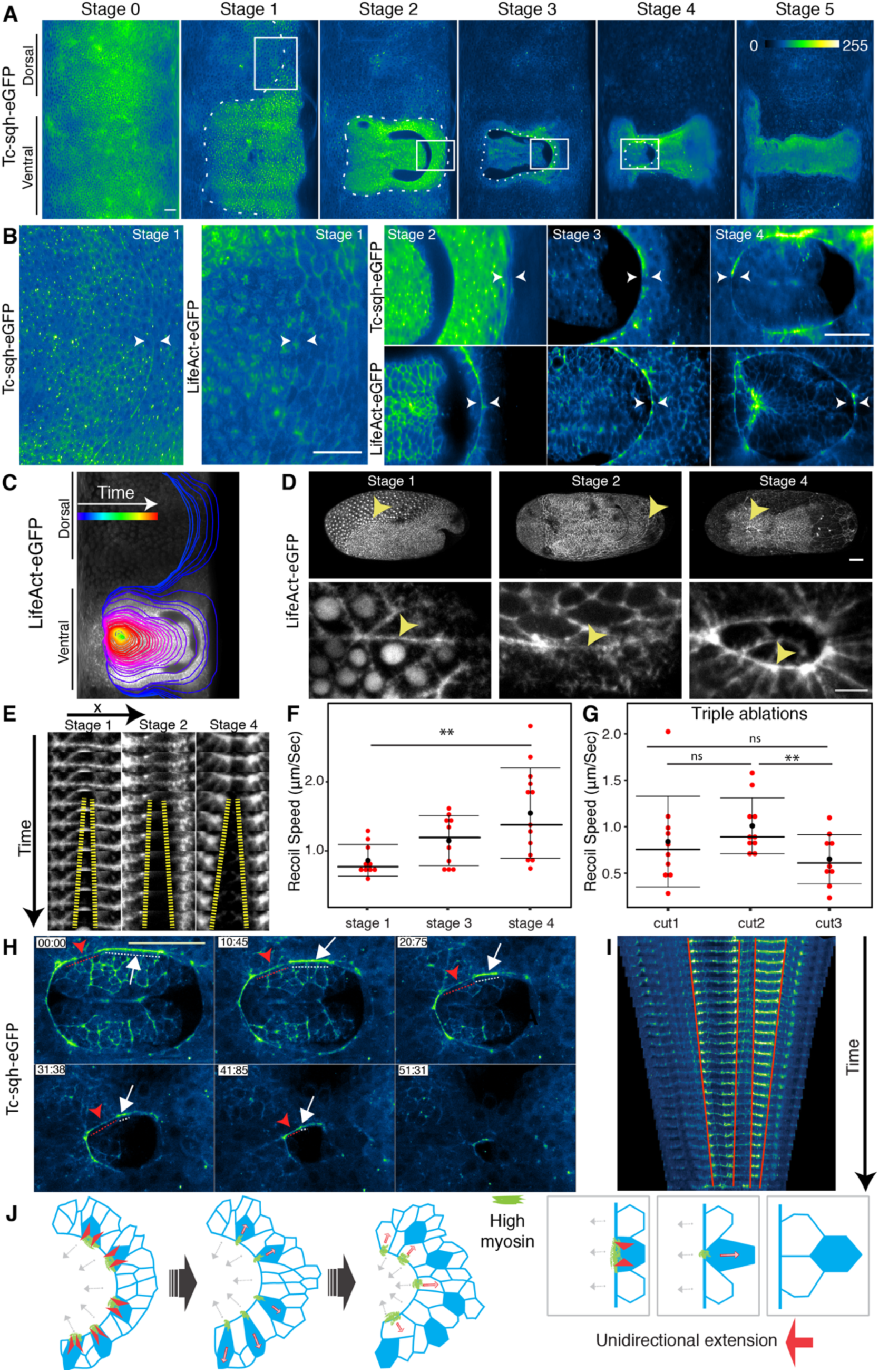
Emergence of a heterogeneous actomyosin cable at the serosa-embryonic boundary promotes cell eviction during serosa window closure. **(A)** Cartographic projection of *Tribolium* embryos injected with Tc-sqh-eGFP and imaged with multi-view SPIM. The accumulation of myosin at the border between serosa and embryos is highlighted by the dotted line as it emerges around the egg circumference (Stage 1) and then during its progressive constriction on the ventral side of the embryo (Stages 2-5). Scale bar is approximately 50 *µ*m. **(B)** Inset shows zoomed in images of Tc-sqh-eGFP localization in the regions marked by white boxes in (A) and images from cartographic projections of an embryo injected with LifeAct-eGFP mRNA and imaged with multi-view SPIM. **(C)** The shape of the supracellular actomyosin cable in map projected LifeAct-eGFP SPIM recording is outlined over time as it emerges dorsally and closes on the ventral side of the embryo. The color of the outline corresponds to the time stamp of the frame from which it was traced. **(D)** Maximum intensity projections of confocal stacks of LifeAct-eGFP injected embryos from 3 different developmental stages. Arrowheads points to the regions of the cable that was ablated. Stage 1 images are lateral views and Stage 2 and 4 images ventral views. Bottom row shows close ups of areas marked by arrows in the top row. Scale bar in top panel is 50 *µ*m and 10 *µ*m in the bottom panels. **(E)** Kymograph of the recoiling membrane edges (yellow hyphen) after laser ablation of the cells forming the actomyosin cable at the leading edge of the serosa window. **(F)** The distributions of recoil velocities after ablation of the cable-forming cells at different stages (n>10). **(G)** The distributions of recoil velocities after three successive laser ablations of three distinct cable-forming cell edges in a single cable at Stage 4 (n >10). Statistical significance determined by paired t-test. **(H)** Images from a timelapse point scanning confocal recording of an embryo expressing Tc-sqh-eGFP. Myosin localization at the cable is different between different cable forming cells. Cell with high myosin accumulation is labelled with white arrow and its extent is highlighted with white dotted line. Cell with low myosin is labelled similarly but in red. Scale bar is 10 *µ*m. **(I)** Kymograph of myosin cable shown in (I). The cable was segmented manually and straightened computationally in Fiji. **(J)** Illustration shows the differential contraction of the serosa-window-facing cell edges depending on the amount of myosin. This leads to T1 transitions in the serosa (right). As a result, the leading edge of serosa extends unidirectionally and at the same time undergoes structural rearrangement. Green color depicts the myosin enriched in the contracting cells (red arrowheads).

Such a model predicts that in the absence of the actomyosin cable the serosa window would fail to close ventrally. Since the actomyosin cable forms at the extraembryonic-embryonic tissue boundary, we hypothesized that we could abolish the emergence of the cable by RNAi knockdown of the *Tribolium* transcription factor-encoding *zerknüllt-1* gene (*Tc-zen1)* that specifies extraembryonic (serosal) cell fate^11^. Live imaging of *Tc-zen1*^*RNAi*^ embryos injected with LifeAct-eGFP revealed indeed the absence of the actomyosin cable (**Fig 5A,B, *Supplementary movie 9***). While such *Tc-zen1*^*RNAi*^ embryos started the contraction and folding of the embryonic primordium as wildtype embryos, the epibolic movement halted and a ventral serosa window failed to form and close (**Fig 5A,B,E, *Supplementary movie 10***). Compared to wildtype, the dorsal spreading cells in *Tc-zen1*^*RNAi*^ embryos became larger, presumably due to their lower number (**Fig 5C**). The cells on the ventral leading edge, however, were much smaller (**Fig 5D,F**), did not elongate anisotropically (**Fig 5B, *Supplementary Fig 3B***), did not exchange neighbors and were not evicted from the leading edge (**Fig 5E**). Finally, although the shape index of the dorsal cells in *Tc-zen1*^*RNAi*^ embryos was comparable to wildtype (**Fig 5G,H)**, the ventral region showed a significantly lower shape index (**Fig 5G,I)** with less pronounced regionalization around the serosal window (**Fig 5G,J**). We conclude that in the absence of the actomyosin cable, the marginal cells fail to become evicted from the leading edge, tissue fluidization fails to occur and, consequently, the epithelial tissue fails to remodel and close its gap.

**Figure 5:**
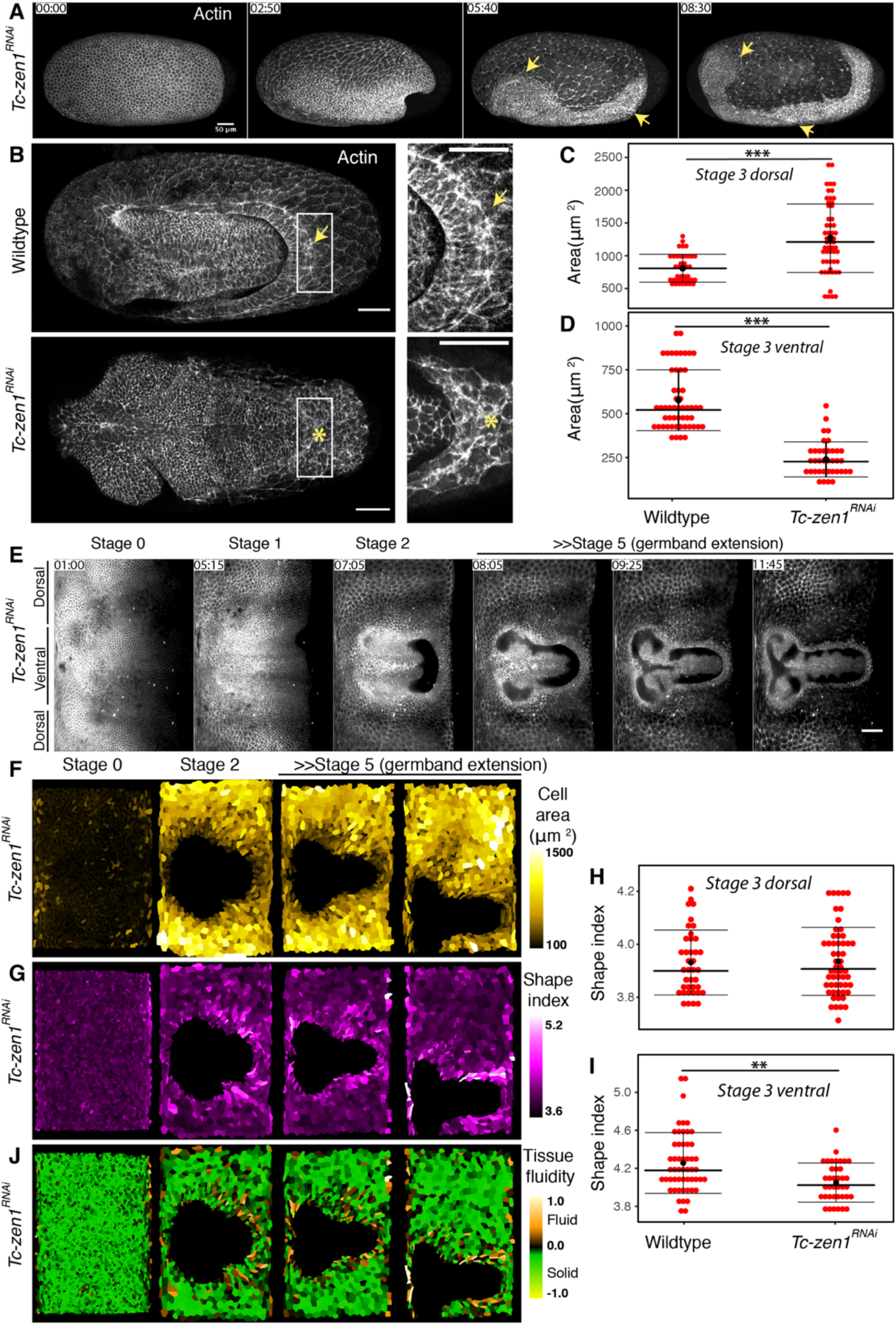
Cell and tissue dynamics in *Tc-zen1* knockdown embryos. **(A)** Maximum intensity projections of a developing embryo labelled with LifeAct-GFP, injected with dsRNA for *Tc-zen1* and imaged with point scanning confocal microscope. Arrows point to the open serosa window in the head and the posterior region. Scale bar is 50 *µ*m. **(B)** Selected maximum intensity projection images from wildtype (top) and *Tc-zen1*^RNAi^ (bottom) embryos at stage 3. Inset shows the cable in wildtype embryos (arrow) and absence of the cable in the knockdown (*). Scale bar is 50 *µ*m. **(C)** The distributions of cell areas in wildtype and *Tc-zen1* knockdown embryos sampled from confocal datasets in the ventral (C) and dorsal (D) serosa regions at Stage 3. The number of cells (n) and the number of embryos (N) sampled at different stages were as follows: In (C) wildtype n=39, N=6, *Tc-zen1*^RNAi^ n=55, N=7, in (D) wildtype n= 52, N=7, *Tc-zen1*^RNAi^ n=38, N=7, **(E)** Cartographic projections of a reconstructed multi-view SPIM recording in which an embryo injected with Gap43-eYFP and *Tc-zen1* dsRNA was imaged from 5 angles every 5 minutes. Scale bar is approximately 100 *µ*m. **(F)** Cartographic projections shown in (E) overlaid with manually curated automated segmentation and colored according to the apical area of the segmented serosa cells. **(G)** Cartographic projections shown in (E) overlaid with manually curated automated segmentation and colored according to the shape index of the segmented serosa cells. **(H)** The distributions of shape indices in wildtype and *Tc-zen1* knockdown embryos sampled from confocal datasets in the dorsal (H) and ventral (**I**) serosa regions at Stage 3. Numbers of cells and embryos are the same as in panel (C). (**J**) Cartographic projections shown in (E) overlaid with manually curated automated segmentation and colored according to the tissue fluidity (see methods) of the segmented serosa cells.

The epibolic expansion of the *Tribolium* serosa to envelop the entire egg surface is a dynamic morphogenetic process constrained by the ellipsoidal geometry of the egg and the mechanical properties of the tissue. Our data suggest that the regionalized tissue fluidization at its leading edge solves the geometrical and mechanical problems associated with serosal epiboly. First, it addresses the geometric constraints necessitating both the expansion and regional compaction of the tissue to close the gap. While the bulk of the tissue expands in a manner similar to an elastic solid material, the fluid-like ventral region remodels, halts the increase in cell area and therefore can remain compact. Second, in the absence of cell divisions, which have been implicated as a stress-release mechanism in fish^4,29^, local cell rearrangements induced by actomyosin contractility at the leading edge release the mechanical stress in the non-proliferative serosal sheet and maintain epithelial integrity during closure.

Our results suggest that the contractile forces of the heterogeneous actomyosin network operate at the single-cell level to exclude marginal cells individually from the serosa window. The order in which cells are evicted is dictated by the local myosin accumulation at each cable-forming edge. This is consistent with previous findings that myosin intensity correlates with tension in wound healing cable^30,31^. Furthermore, it has been suggested that a non-uniform stepwise contractility of individual edges is necessary for efficient epithelial closure during wound healing in *Drosophila* embryos and neural tube closure in chordates^32,33^. This kind of sequential contraction is likely operating during window closure to dissipate serosa resistance. Last but not the least, a recent study proposed that tissue fluidization is required for seamless wound healing in damaged *Drosophila* imaginal discs^18^. Similar to the actomyosin cable of the *Tribolium* serosa window, the cable that assembles at the leading edge of the wound evicts cells from the wound periphery and promotes cell intercalation resulting in tissue fluidization and acceleration of epithelial gap closure. All these striking similarities point towards a conserved morphogenetic function of actomyosin cables in shaping and repairing epithelia by local tissue fluidization.

## Acknowledgements

We would like to thank Thorsten Horn for teaching us various *Tribolium* techniques, Matthew A. Benton for kindly providing the pCS2+LifeAct-eGFP plasmid and sharing embryonic injection protocol and discussions, Sebastian Streichan for optimising ImSANE for *Tribolium* SPIM data and critical discussions, Alexander Dibrov for helping with tissue cartography cell segmentations, Matthias Merkel for providing code for tissue shape analysis, Christopher Schmied for optimizing Snakemake SPIM data analysis pipeline for our datasets, Michaela Burkon for helping with *Tribolium* stock keeping and in doing parental RNAi experiments, MPI-CBG Light Microscopy Facility for help with imaging, Mette Handberg-Thorsager and Yu-Wen Hsieh for sharing plasmids and help with cloning, Ivana Viktorinova for schematic drawings, Anna Giles and Johannes Schinko (Averof lab), Peter Kitzmann (Bucher lab) and the van der Zee lab for sharing valuable transgenic lines and Siegfried Roth for critical discussions.

## Author Contributions

A.J. designed the research, performed experiments, analyzed the data, and wrote the manuscript. V.U produced image analysis software and analyzed data. A.M contributed to data analysis, M.P helped in segmenting data, L.P helped in laser ablation experiments, S.M contributed reagents, data and was involved in discussions, R.H. produced image analysis software and contributed to analysis workflow design, K.A.P. conducted RNAi parameter validation experiments and was involved in discussions, F.J. contributed to data segmentation, S.W.G helped in interpreting laser ablation data and was involved in discussions, P.T. and A.P. conceived and oversaw the project, and wrote the manuscript.

## Methods

### *Tribolium* rearing and stocks

*Tribolium castaneum* stocks were kept at 32°C and 70% relative humidity on whole-grain or white flour supplemented with yeast powder according to standard procedures^34^. All mRNA injections were performed into embryos of the *vermilion*^*white*^ strain. The following transgenic lines were used for live imaging: i) EFA-nGFP, ubiquitously expressing a nuclear-localized GFP reporter^35^ (kindly provided by Michalis Averof’s lab); ii) αTub-H2A-eGFP, ubiquitously expressing a nuclear eGFP reporter (kindly provided by Peter Kitzmann from Gregor Bucher’s lab); EFA-Gap43-YFP,2A-Histone-RFP, ubiquitously expressing both a membrane YFP and a nuclear RFP reporter (kindly provided by Johannes Schinko and Anna Gilles from Michalis Averof’s lab); iv) αTub-LifeAct-eGFP, ubiquitously labelling filamentous actin with eGFP^16^ (kindly provided by the Van der Zee lab); v) αTub-Tc-sqh-eGFP, ubiquitously labelling the *Tribolium* non-muscle myosin II through its regulatory light chain (Tc-sqh). Details about the αTub-Tc-sqh-eGFP transgenesis construct are available upon request. Overview of genotypes and constructs used in the study is provided in ***Supplementary Methods Table 1.***

### RNA injections

Actin and myosin dynamics were visualized in *vermilion*^*white*^ embryos injected with *in vitro* transcribed capped mRNAs encoding LifeAct-eGFP or Tc-sqh-eGFP that were synthesized from linearized plasmid templates pT7-LifeAct-eGFP and pCS2+-Tc-sqh-eGFP, respectively^13,28^. For the RNAi knock-down experiments of *Tc-zen1*, the dsRNA against the *Tribolium zerknüllt-1* gene (TC000921) was synthesized with primers optimized for gene specificity (a 203-bp amplicon outside of the conserved homeobox region)^36^. The mRNAs and the dsRNA were each injected at a final concentration of 1 mg/ml. Eggs from the *vermilion*^*white*^ strain were collected for two hours at 30°C, aged for another hour at 30°C and dechorionated in 16% commercial Klorix bleach for 1 to 2 minutes. Dechorionated pre-blastoderm embryos were mounted on a 1% agar bed and were microinjected in air through their anterior pole under a brightfield upright microscope as previously described^13,34^(***Supplementary Fig 1A***). Injected eggs were incubated in humid chambers at 30°C for ∼ 2 hours and the most homogeneously labeled and bright embryos were selected for imaging. For parental knock-down of *Tc-zen1* by RNAi, dsRNA was injected into the abdomen of female pupae collected from the αTub-LifeAct-eGFP transgenic line. Injected adult females were crossed to males from the same line and their eggs were collected for imaging.

### Live imaging with confocal and light-sheet microscopy

Confocal live imaging was carried out at 25°C or 30°C on an inverted Zeiss LSM 780 system equipped with a temperature-controlled incubator. Embryos were mounted in 1% agarose in glass bottom petri dishes and covered in water. Embryos were scanned with a Zeiss 25x/0.8 NA Plan-Apochromat multi-immersion objective or a Zeiss 40x/1.2 NA C-Apochromat water-dipping objective with pixel sizes ranging between 0.2 µm and 0.55 µm, a z-step of 2 μm and a temporal resolution of 5 minutes. Multi-view light-sheet imaging (referred to as Selective Plane Illumination Microscopy (SPIM) or light sheet microscopy) was carried out on a Zeiss Lightsheet Z.1 microscope equipped with a 20x/1.0 NA Plan Apochromat water-immersion detection objective and two 10x/0.2 NA dry illumination objectives. Embryos were embedded in glass capillaries in 1% low melting agarose dissolved in 1xPBS together with fluorescent beads as previously describe^37,38^. For each embryo, z-stacks were acquired from 5 views every 72° with voxel size 0.381 µm × 0.381 µm × 2.0 µm. The starting point in the time-stamps used for all experiments was the last (12^th^) round of synchronous nuclear divisions which precedes the formation of the uniform blastoderm and all subsequent morphogenetic events^12^. Parameters for all live imaging experiments are summarized in the ***Supplementary Methods Table 1***.

### Laser ablations

Laser ablations were performed either on an inverted Zeiss LSM 780 NLO with a 40x/1.2 NA water-dipping objective using an 800nm pulsed infrared laser or on a customized spinning disc confocal unit with a 63x water-dipping objective using an ultraviolet laser microdissection apparatus similar to the one described in^39^. On the first system, three planes with 1-2µm z-spacing were imaged every 1.6 sec (Fig 4F), 2.5 sec (Fig 4G) 2.6 sec (Fig 3C) and the cut was performed in the middle plane, while on the latter system a single plane was recorded every 0.5 second (Fig 3D). Tissue cuts were about 12 µm long spanning 3 to 4 cell diameters, while ablations of single edges were about 5 µm long. The recoil velocity of ablated edges was measured between 6 post-cut time frames using the manual tracking plugin in Fiji. For each cut, two to three independent tracks of the recoiling tissue edges were averaged. The initial recoil velocity was estimated using standard fitting procedures^40^.

### Image processing

The multi-view light-sheet datasets were registered and fused using Fiji plugins as previously described^41–43^. The 4D (3D+time) fused datasets were converted into 3D (2D+time) time-lapse maps by making cylindrical projections using the ImSANE software^15^. Cells were segmented using a deep learning-based approach called StarDist which is capable of learning morphological priors^44^. Different neural networks were trained for different markers (membrane and actin labels). The training data were obtained by generating realistic looking synthetic microscopic images of Tribolium using Generative Adversarial Networks (GANs). The generated synthetic data were evaluated visually against the real microscopy data to ensure textural and morphological consistency between the two. After training StarDist networks on such synthetic data, they were applied to the real microscopy images and the predictions were manually curated in Labkit (Fiji plugin, http://sites.imagej.net/Labkit/) to fix any segmentation mistakes. After cartographic projections, some cells on the horizontal (top/bottom) edges of the maps were necessarily cut in order to unfurl the 3D embryo to 2D. Those incomplete cells were excluded from analysis. Distortions that are inherent to the mapping of curved surfaces onto a plane were corrected with custom Fiji plugins (available on the “Tomancak lab” Fiji Update site) thereby allowing the measurement of quantities like size, circularity, shape factor, density, velocity, and the local cell alignment (see below). Consequently, the scale bars in map projections are only approximate and reflect accurately the sizes only in the middle portions of the maps. Nuclei in the depth color-coded cartographic projections were tracked using MaMuT^38^ and Mastodon (both available via Fiji Update sites).

### Shape index analysis

Shape index was calculated for each segmented cell in the 2D cartographic projections as ***p*** = **P**/√(**A**) where **P** is the cell perimeter and **A** is the cross-sectional area^22^. The measurements were distortion-corrected using the above-mentioned Fiji plugins and plotted onto the segmented cartographic projection as a color map.

The local *cell shape alignment index* **Q** was calculated as in Wang et al., 2019^25^. Briefly, cells in the map projection were converted into a triangular mesh connecting the centers of all adjacent segmented cells. That is, where three (or more) cells touch, a triangle (or a triangle fan) with vertices coinciding with centers of adjacent cells was formed. For every triangle, a degree *q* of deviation from equilateral triangle was computed^27^. For every cell then, its shape alignment index **Q** became a weighted average over *q* from all triangles whose vertex coincides with this cell’s center. Using this **Q**, an adjusted shape index threshold was determined as *p*_*adj*_= *p*_*o*_ + 4*bQ*^2^ for *p*_*o*_=3.94 and *b*=0.43^25^. According to Wang et al. 2019 simulations this threshold marks solid-to-fluid transition for a given anisotropy in the tissue (i.e. for a given value of cell shape alignment index **Q**). The tissue fluidity for a given cell was then calculated as a difference between its actual shape index *p* and the *p*_*adj*_ for a given local value of **Q**. This difference was converted into a color code and displayed on the cartographic projection. Green color signifies solid-like local tissue properties (***p*** < *p*_*adj*_), brown color fluid-like local tissue properties (***p*** > *p*_*adj*_) and black color marks the vicinity to the theoretically predicted solid-to-fluid transition (***p*** = *p*_*adj*_).

---

**Supplementary Figure 1:**
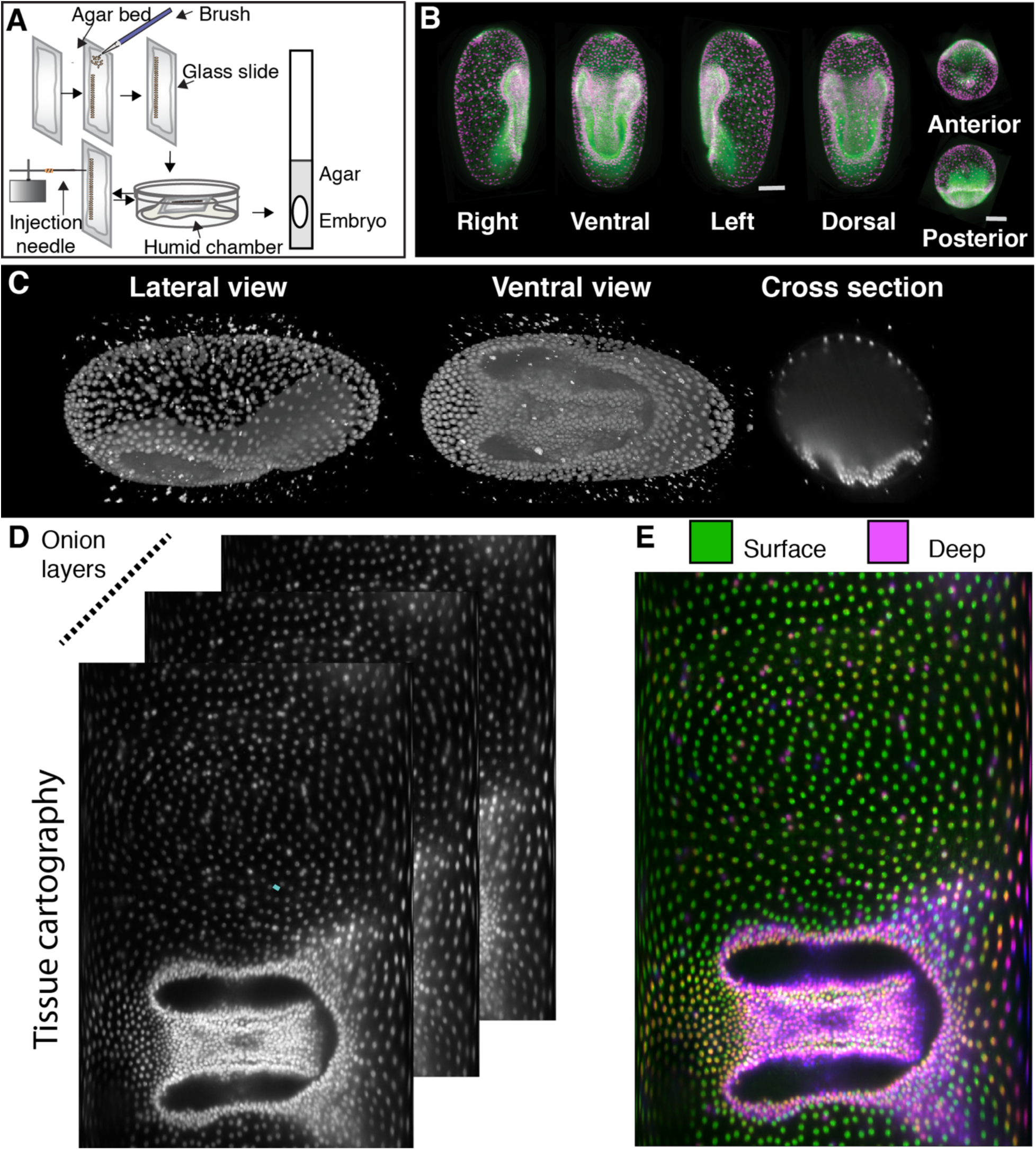
*Tribolium* embryo preparation, imaging and image analysis pipeline to study serosa epiboly. **(A)** Illustration outlines the micro-injection and sample mounting protocol to label *Tribolium* embryos and mount them for lightsheet microscopy. **(B)** Maximum intensity projections from different orientation of 3D images of embryos injected with LifeAct-eGFP and Histone-RFP. The embryo was imaged from 5 views with a light sheet microscope. Individual view stacks were registered and fused based on fluorescent beads scattered in the mounting medium (bright dots surrounding embryo) using Fiji Multi-view Reconstruction plugin. **(C)** 3D rendering of Histone-eGFP expressing *Tribolium* embryo reconstructed from multi-view SPIM data and viewed from the ventral and lateral side. Sagittal cross section of the same embryo volume (right). **(D)** The embryo shown in C is dimensionality reduced from 3D to 2D by generating a cartographic projection. Successive, increasingly deep concentric layers of the embryo surface are shown as separate maximum intensity projections. **(E)** The different layers generated in (D) are color coded to separate the superficial serosa from the internalized embryo.

**Supplementary Figure 2:**
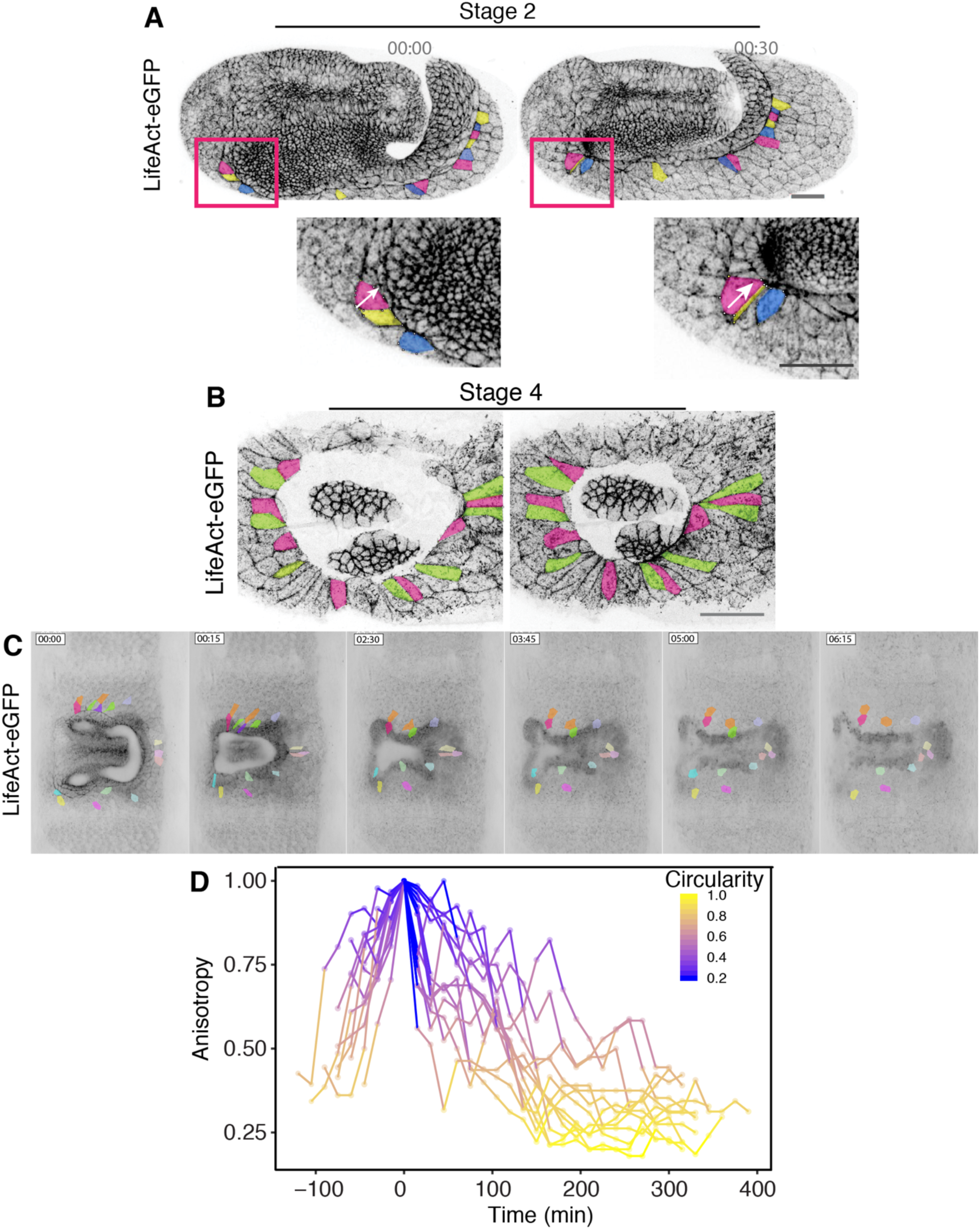
Anisotropy of the cells at the leading edge of the serosa window increases over time and decreases once they leave the edge. **(A)** Inverted confocal images of a Life-Act-eGFP expressing *Tribolium* embryos show selected cell outlines (highlighted in different colors) at the leading edge of the serosa window at Stage 2. Insets zoom in on regions highlighted with red boxes. Arrow points to a cell elongating along the axis roughly perpendicular to the leading edge. Scale bar is 50 *µ*m. **(B)** Inverted confocal images similar to (A) of LifeAct-eGFP expressing *Tribolium* embryos show selected elongated cell outlines at the leading edge of the serosa window at Stage 4. Scale bar is 10 *µ*m. **(C)** Inverted cartographic projections of LifeAct-eGFP expressing *Tribolium* embryos imaged with multi-view SPIM show outlines of selected anisotropically elongated cells during serosa window closure. The highlighted cells increase their shape anisotropy over time till they leave the leading edge of the serosa window and become hexagonal. **(D)** Graph shows the change in anisotropy of cells highlighted in (C) over time. Anisotropy is defined as deviation from the circle that has circularity value of 1.

**Supplementary Figure 3:**
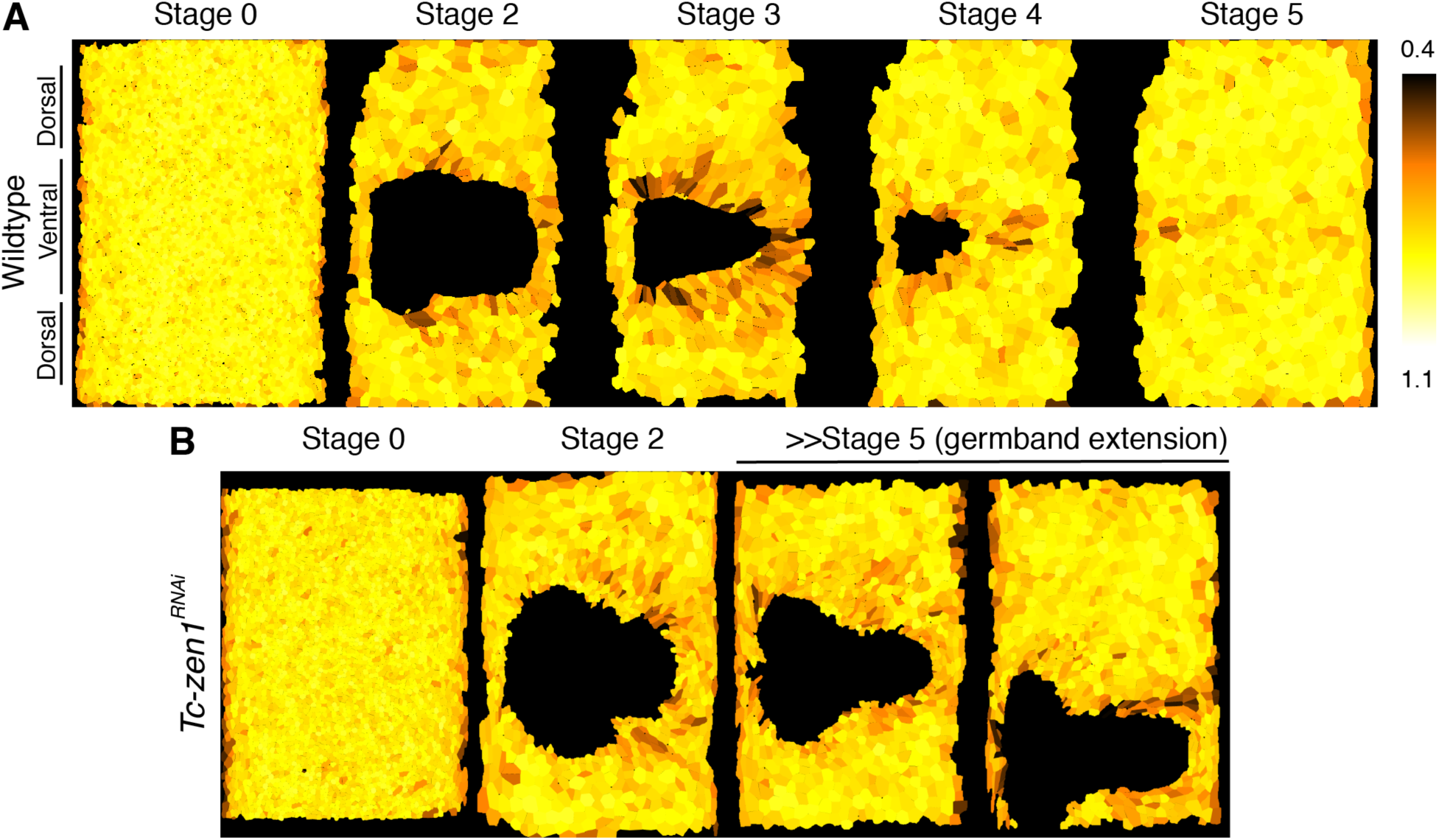
Cell shape anisotropy in wildtype and *Tc-zen1*^*RNAi*^ embryos. **(A)** Cartographic projections of reference stages of wildtype embryo labelled with LifeAct-eGFP and imaged live with multi-view SPIM. The projections are overlaid with manually curated automated segmentation results visualizing anisotropy of serosa cells through a color code. Anisotropy is defined as deviation from the circle that has circularity value of 1. **(B)** Cartographic projections of a multi-view SPIM recording in which embryos injected with Gap43-eYFP and *Tc-zen1* dsRNA was imaged from 5 angles every 5 minutes. The projections are color coded as in (A).

**Supplementary Figure 4:**
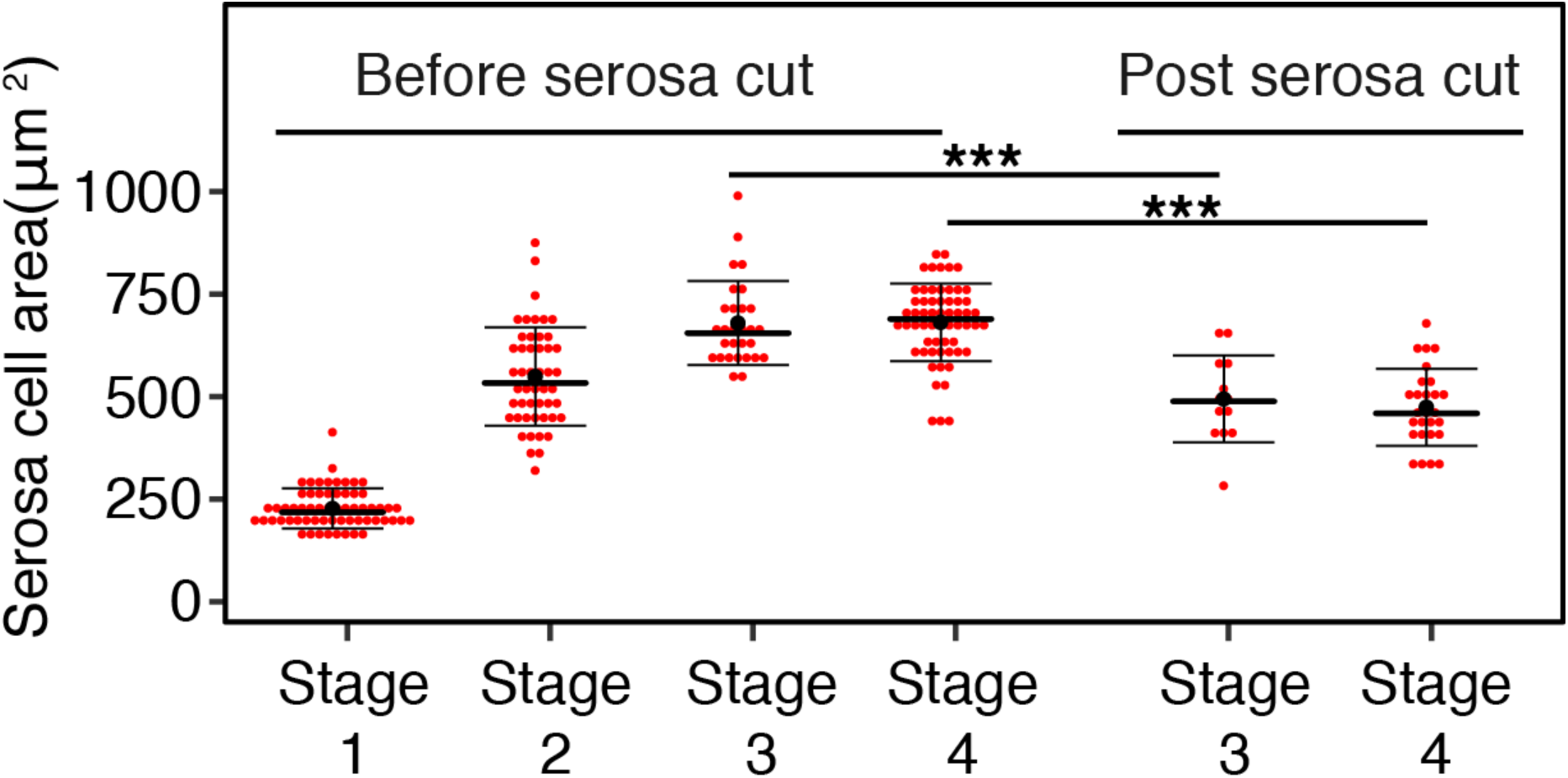
Contraction of cells after release in serosal tension. Graph showing apical cell areas in the serosa before and after laser ablations at different reference stages. Intact cells neighboring the ablation site in embryos expressing LifeAct-eGFP were measured before and after laser cuts in the dorsal serosa.

**Supplementary Figure 5:**
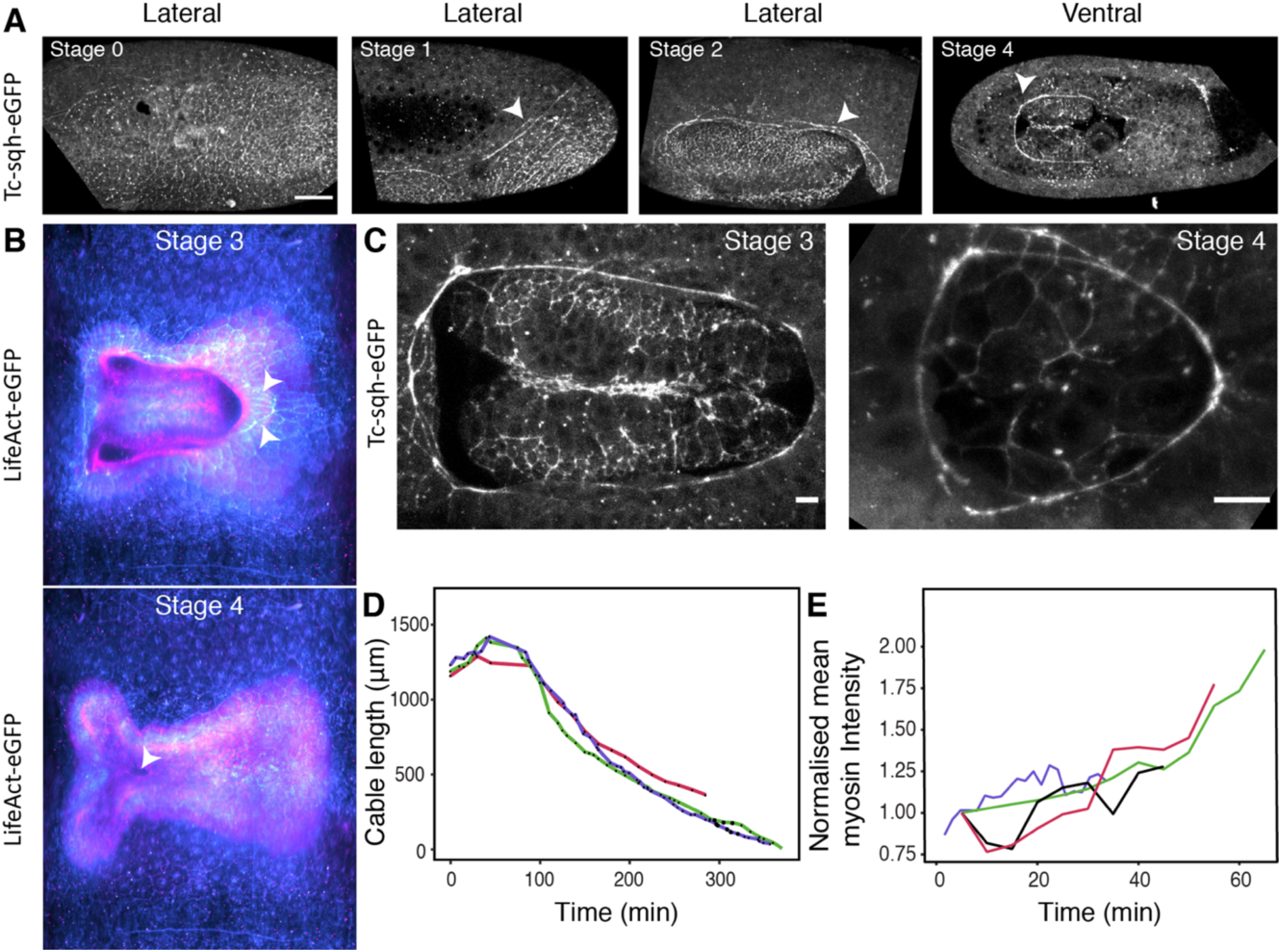
Actomyosin localization at the embryo-serosa boundary. **A**) Myosin localization in *Tribolium* embryo at different stages of gastrulation morphogenesis. Arrowheads point to the myosin cable. The embryos were injected with mRNA encoding Tc-sqh-eGFP and imaged with point scanning confocal microscope. Scale bar is 50 *µ*m. **(B)** Images show the ventral region of a stage 3 and 4 embryo from a cartographic projection. The embryo was labeled with LifeAct-eGFP and imaged with multi-view SPIM. The different layers of the cartographic projection are colored with cyan for surface serosa and magenta for internal embryo layers. The arrowheads point to the actin cable at the serosa window leading edge at Stage 3. **(C)** Confocal images show myosin enrichment at the embryo and serosa boundary at stage 3 and stage 4. Scale bar is 10 *µ*m. **(D)** The graph shows the length of manually segmented actomyosin cable as a function of time during serosa window closure (N=3). **(E)** The graph shows mean myosin intensity normalized to the initial value at the manually segmented cable over time (N=4).

## Supplementary movies

**Supplementary Movie 1:** Lateral and ventral views of 3D rendered multi-view Lightsheet recording of *Tribolium* embryo collected from a Histone-eGFP transgenic line. The embryo was imaged from 5 angles at 1.5 minute time interval at 22°C. Time stamp is hh:mm.

**Supplementary Movie 2:** The expanding serosa is outlined on a 2D cartographic projection of a 4D SPIM recording of a transgenic embryo from the EFA-nGFP line marking cell nuclei. The color of the serosa changes according to its increasing total area. Time stamp is hh:mm.

**Supplementary Movie 3:** 4D multi-view lightsheet recording of a *Tribolium* embryo expressing EFA-nGFP nuclear marker projected as a 2D cartographic map. The embryo was imaged from 5 angles at 1.5 minutes time interval. Successive onion layers of the map are color coded to distinguish deeper embryo layers from the superficial serosa. The last few cells contributing to the serosa window closure are tracked from Stage 1 onwards. Tracks are color-coded by time. Time stamp is hh:mm.

**Supplementary Movie 4:** Timelapse video shows Stage 4 to serosa window closure in a LifeAct-GFP labelled embryo imaged with point scanning confocal microscope. Selected cells at the cable are highlighted and tracked. Time stamp is hh:mm. Scale bar is 50 *µ*m.

**Supplementary Movie 5:** *Tribolium* embryo labelled with LifeAct-eGFP was imaged using 4D multi-view lightsheet microscopy and projected as a cartographic map. Dots and lines show two groups of serosal cells on the dorsal and ventral side of the embryo tracked over time using Mastodon Fiji Plugin. Time stamp is hh:mm.

**Supplementary Movie 6:** Cartographic projection of 4D multi-view lightsheet recording of an embryo injected with Tc-sqh-eGFP at 22°C. The dorsal part of the embryo is positioned in the middle to show the emergence of the myosin cable at Stage 1 (pointed out by an arrow). Time stamp is hh:mm.

**Supplementary Movie 7:** Timelapse videos of Stage 1, 3 and 4 embryos labelled with LifeAct-eGFP. The edges of serosa leading edge cells facing the serosa window (where cable-like actin enrichment occurs) were laser ablated and the edges were tracked with Fiji to measure the recoil velocity over time. Time stamp is mm:ss.

**Supplementary Movie 8:** Timelapse video of a Stage 4 transgenic embryo labelled with Tc-sqh-eGFP and imaged with a point scanning confocal microscope. The 3D stacks were maximum intensity projected. Myosin is distributed in a heterogeneous manner along the cable with different cell edges showing different intensities (highlighted using Green-Blue look-up table; green is high myosin). Time stamp is mm:ss.

**Supplementary Movie 9:** Timelapse video of an embryo labelled with LifeAct-eGFP in which *Tc-zen1* was knocked down using parental RNAi. The embryo was imaged with a point scanning confocal microscope and the 3D stacks were maximum intensity projected. Time stamp is hh:mm.

**Supplementary Movie 10:** Cartographic projection of a multi-view lightsheet dataset. The embryo was injected with mRNA for GAP43-eYFP to label cell membranes and dsRNA to knockdown *Tc-zen1*. Time stamp is hh:mm.

**Supplementary Movie 11:** Cartographic maps of serosal areas (Fig 1G), shape index (Fig 2G), circularity (Supplementary Fig 3), fluidity (Fig 3E) back projected to the original 3D volume and volumetric rendered them using Fiji.

## Supplementary Methods Table

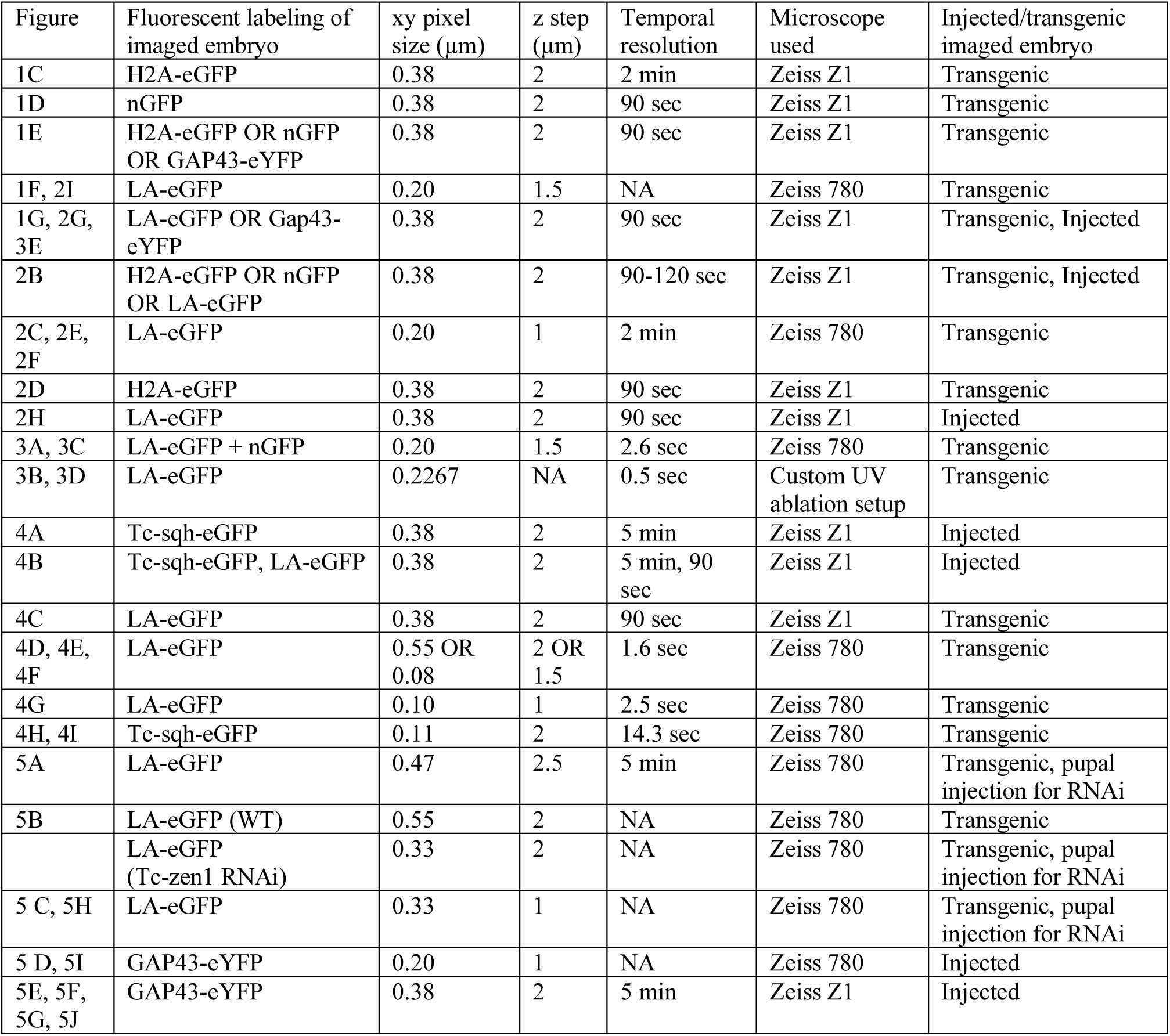

## Notes

#### Summary of Updates

Thoroughly revised and updated manuscript.

## References

1. Solnica-Krezel, L. Conserved patterns of cell movements during vertebrate gastrulation. Curr. Biol. 15, R213–28 (2005).

2. Keller, R. E. & Trinkaus, J. P. Rearrangement of Enveloping Layer Cells Without Disruption of the Epithelial Permeability Barrier as a Factor in Fundulus Epiboly. Dev. Biol. 120, 12–24 (1987).

3. Köppen, M. Coordinated cell-shape changes control epithelial movement in zebrafish and Drosophila. Development 133, 2671–2681 (2006).

4. Campinho, P. et al. Tension-oriented cell divisions limit anisotropic tissue tension in epithelial spreading during zebrafish epiboly. Nat. Cell Biol. 15, 1405–1414 (2013).

5. Behrndt, M. et al. Forces Driving Epithelial Spreading in Zebrafish Gastrulation. Science 338, 257–260 (2012).

6. Panfilio, K. A. Extraembryonic development in insects and the acrobatics of blastokinesis. Dev. Biol. 313, 471–491 (2008).

7. Schmidt-Ott, U. & Kwan, C. W. Morphogenetic functions of extraembryonic membranes in insects. Curr. Opin. Insect Sci. 13, 86–92 (2016).

8. Jacobs, C. G. C. & van der Zee, M. Immune competence in insect eggs depends on the extraembryonic serosa. Dev. Comp. Immunol. 41, 263–269 (2013).

9. Jacobs, C. G. C., Rezende, G. L., Lamers, G. E. M. & van der Zee, M. The extraembryonic serosa protects the insect egg against desiccation. Proc. R. Soc. B Biol. Sci. 280, 20131082–20131082 (2013).

10. Caroti, F. et al. Decoupling from yolk sac is required for extraembryonic tissue spreading in the scuttle fly Megaselia abdita. eLife 7, e34616 (2018).

11. van der Zee, M., Berns, N. & Roth, S. Distinct Functions of the Tribolium zerknüllt Genes in Serosa Specification and Dorsal Closure. Curr. Biol. 15, 624–636 (2005).

12. Handel, K., Grünfelder, C. G., Roth, S. & Sander, K. Tribolium embryogenesis: a SEM study of cell shapes and movements from blastoderm to serosal closure. Dev. Genes Evol. 210, 167–179 (2000).

13. Benton, M. A., Akam, M. & Pavlopoulos, A. Cell and tissue dynamics during Tribolium embryogenesis revealed by versatile fluorescence labeling approaches. Development 140, 3210–3220 (2013).

14. Benton, M. A. & Pavlopoulos, A. Tribolium embryo morphogenesis: may the force be with you. BioArchitecture 4, 16–21 (2014).

15. Heemskerk, I. & Streichan, S. J. Tissue cartography: compressing bio-image data by dimensional reduction. Nat. Methods 12, 1139–1142 (2015).

16. van Drongelen, R., Vazquez-Faci, T., Huijben, T. A. P. M., van der Zee, M. & Idema, T. Mechanics of epithelial tissue formation. J. Theor. Biol. 454, 182–189 (2018).

17. Benton, M. A. et al. Fog signaling has diverse roles in epithelial morphogenesis in insects. eLife 8, (2019).

18. Tetley, R. J. et al. Tissue fluidity promotes epithelial wound healing. Nat. Phys. 15, 1195–1203 (2019).

19. Mongera, A. et al. A fluid-to-solid jamming transition underlies vertebrate body axis elongation. Nature 561, 401–405 (2018).

20. Lawton, A. K. et al. Regulated tissue fluidity steers zebrafish body elongation. Development 140, 573–582 (2013).

21. Petridou, N. I., Grigolon, S., Salbreux, G., Hannezo, E. & Heisenberg, C.-P. Fluidization-mediated tissue spreading by mitotic cell rounding and non-canonical Wnt signalling. Nat. Cell Biol. 21, 169–178 (2019).

22. Bi, D., Lopez, J. H., Schwarz, J. M. & Manning, M. L. A density-independent rigidity transition in biological tissues. Nat. Phys. 11, 1074–1079 (2015).

23. Bi, D., Yang, X., Marchetti, M. C. & Manning, M. L. Motility-Driven Glass and Jamming Transitions in Biological Tissues. Phys. Rev. X 6, 021011 (2016).

24. Yang, X. et al. Correlating cell shape and cellular stress in motile confluent tissues. Proc. Natl. Acad. Sci. 114, 12663–12668 (2017).

25. Wang, X. et al. Anisotropy links cell shapes to a solid-to-fluid transition during convergent extension. http://biorxiv.org/lookup/doi/10.1101/781492 (2019).

26. Smutny, M., Behrndt, M., Campinho, P., Ruprecht, V. & Heisenberg, C.-P. UV laser ablation to measure cell and tissue-generated forces in the zebrafish embryo in vivo and ex vivo. Methods Mol. Biol. 1189 219–235 (Springer New York, 2014).

27. Merkel, M. et al. Triangles bridge the scales: Quantifying cellular contributions to tissue deformation. Phys. Rev. E 95, 032401 (2017).

28. Münster, S. et al. Attachment of the blastoderm to the vitelline envelope affects gastrulation of insects. Nature 568, 395–399 (2019).

29. Firmino, J., Rocancourt, D., Saadaoui, M., Moreau, C. & Gros, J. Cell Division Drives Epithelial Cell Rearrangements during Gastrulation in Chick. Dev. Cell 36, 249–261 (2016).

30. Fernandez-Gonzalez, R., Simoes, S. de M., Röper, J.-C., Eaton, S. & Zallen, J. A. Myosin II Dynamics Are Regulated by Tension in Intercalating Cells. Dev. Cell 17, 736–743 (2009).

31. Kobb, A. B. & Zulueta-Coarasa, T. Tension regulates myosin dynamics during Drosophila embryonic wound repair. J. of Cell Sci. 130, 689–696 (2017).

32. Zulueta-Coarasa, T. & Fernandez-Gonzalez, R. Dynamic force patterns promote collective cell movements during embryonic wound repair. Nat. Phys. 14, 750–758 (2018).

33. Hashimoto, H., Robin, F. B., Sherrard, K. M. & Munro, E. M. Sequential Contraction and Exchange of Apical Junctions Drives Zippering and Neural Tube Closure in a Simple Chordate. Dev. Cell 32, 241–255 (2015).

34. Brown, S. J. et al. The Red Flour Beetle, Tribolium castaneum (Coleoptera): A Model for Studies of Development and Pest Biology. Cold Spring Harbor Protocols 8, pdb.emo126 (2009).

35. Sarrazin, A. F., Peel, A. D. & Averof, M. A Segmentation Clock with Two-Segment Periodicity in Insects. Science 336, 338–341 (2012).

36. Gurska, D., Jentzsch, I. & Panfilio K.A. Mutual regulation underlies paralogue functional diversification. biorxiv.org. doi:10.1101/427245, (2018).

37. Schmied, C. & Tomancak, P. Sample Preparation and Mounting of Drosophila Embryos for Multiview Light Sheet Microscopy. Methods Mol. Biol. Clifton NJ 1478, 189–202 (2016).

38. Wolff, C. et al. Multi-view light-sheet imaging and tracking with the MaMuT software reveals the cell lineage of a direct developing arthropod limb. Elife 7, 375 (2018).

39. Grill, S. W., Gönczy, P., Stelzer, E. H. K. & Hyman, A. A. Polarity controls forces governing asymmetric spindle positioning in the Caenorhabditis elegans embryo. Nature 409, 630–633 (2001).

40. Mayer, M., Depken, M., Bois, J. S., Jülicher, F. & Grill, S. W. Anisotropies in cortical tension reveal the physical basis of polarizing cortical flows. Nature 467, 617–621 (2010).

41. Preibisch, S., Saalfeld, S., Schindelin, J. & Tomancak, P. Software for bead-based registration of selective plane illumination microscopy data. Nat. Methods 7, 418–419 (2010).

42. Schindelin, J. et al. Fiji: an open-source platform for biological-image analysis. Nat. Methods 9, 676–682 (2012).

43. Schmied, C., Steinbach, P., Pietzsch, T., Preibisch, S. & Tomancak, P. An automated workflow for parallel processing of large multiview SPIM recordings. Bioinformatics 32, 1112–1114 (2016).

44. Schmidt, U., Weigert, M., Broaddus, C. & Myers, G. Cell Detection with Star-Convex Polygons. in Medical Image Computing and Computer Assisted Intervention 11071 265–273 (Springer International Publishing, 2018).

